# Optimized methods for measuring extracellular ATP from human airway epithelial cells and bronchoalveolar lavage fluid

**DOI:** 10.64898/2026.02.03.703573

**Authors:** Ryan Singer, Elena Kum, Quynh Cao, Jenny P. Nguyen, Wafa Hassan, Suzanne Beaudin, Imran Satia, Jeremy A. Hirota

## Abstract

Extracellular adenosine triphosphate (eATP) is a mediator of purinergic signalling in the airways, implicated in mucociliary function, inflammation, and cough via activation of P2X3 receptors. Elevated airway eATP has been associated with multiple respiratory diseases, yet reliable measurement of eATP remains challenging due to its rapid enzymatic degradation and confounding contributions from intracellular ATP. Here, we describe an optimized, microwell plate-based luminescence method for quantifying eATP from human airway epithelial cell cultures and bronchoalveolar lavage (BAL) fluid with enhanced signal stability. Using a commercially available ATP detection assay with a prolonged luminescence half-life, we introduced a simple 0.45 µm syringe filtration step to remove cells and thereby isolate extracellular ATP. This approach demonstrated ATP specificity via apyrase degradation, and provided a linear detection range from 5 nM to 5 µM. Addition of ATP stabilization buffer preserved ATP levels in cell culture media for at least 4 hours at 4 °C and in human BAL samples for at least 6 weeks at −80°C. Applying this method to primary human bronchial epithelial cells revealed detectable eATP release, with preferential secretion at the apical surface under air-liquid interface conditions. Collectively, this optimized assay enables robust, high-throughput, and time-flexible quantification of eATP in both experimental and clinical airway samples. These methods support improved investigation of purinergic signalling in airway health and disease and may facilitate biomarker development relevant to eATP in the airways.

## Introduction

Extracellular adenosine triphosphate (eATP) is a ubiquitous signalling molecule that is released by stressed or damaged cells that can activate a range of purinergic receptors capable of driving inflammation, immune activation, and also evoking coughing by binding to P2X3 receptors on vagal c-fibres.[1–3] High eATP concentrations in human airways have been found to correlate with COPD severity, asthma, and CF-associated neutrophilic inflammation, suggesting dysfunctional ATP release and/or metabolism in disease states.[4–7] Patients with refractory chronic cough (RCC) also demonstrate an exaggerated cough response to ATP inhalation, suggesting eATP in the airways plays a role in evoking cough.[8] Indeed, gefapixant is an oral P2X3 receptor antagonist approved in Europe, UK, and Japan while camlipixant, another oral P2X3 receptor antagonist, is in phase 3 clinical development.[9, 10] Measuring eATP in the airways has the potential to be used as a potential diagnostic or predictive biomarker guiding P2X3 receptors antagonist administration, but measure of this signaling molecule has been challenging as it is rapidly broken down by ectonucleotide enzymes.

Measurement of eATP *in vitro* is typically performed in real time via transient chemiluminescence reactions using firefly luciferase, which emits light proportionally to ATP concentration within 5-10,000 nM ATP.[11, 12] The half-life of these “flash-type” signals is typically below 30 seconds, limiting the potential for higher throughput or time-delayed experimental designs. Microwell plate-based “glow-type” ATP detection methods in which ATP is quantified from cell culture media samples have longer half-lives ranging from 1-60 minutes depending on the supplier, and are typically not specific for eATP as they lyse cells to release intracellular ATP.

This work describes the development of a microwell plate-based method for measuring eATP from *in vitro* human airway epithelial cell cultures and human bronchoalveolar lavage (BAL) fluids with high signal stability, allowing for more rigorous experimental designs for the analysis of ATP release dynamics. In this approach, an existing luminescence detection assay system with a half-life of 5 hours is modified using small-pore syringe filters to isolate an eATP signal.

## Methods

### Method characterization

The ATPlite™ Luminescence ATP Detection Assay System (Revvity, MA, USA) was used to measure eATP from BAL fluids, cell suspensions, cell culture media, and apical washes of primary human bronchial epithelial cell cultures. The ATPlite™ kit was chosen as it has the unique advantage of providing high ATP signal stability (> 5 h half-life) via an ATP stabilization buffer that irreversibly inhibits ATPases. However, the kit is intended for measuring total ATP levels in cell cultures, including intracellular ATP, via cell lysis reagents contained within the ATP stabilization buffer. Our signal of interest is the active release of ATP into the extracellular space, which would be confounded by cell lysis and subsequent measurement of intracellular ATP. We hypothesized that removing cells from collected cell culture media or human BAL samples would allow for the detection of only eATP with high signal stability. **Figure 1A** displays a graphic flow chart of the ATPlite™ assay procedure according to the manufacturer’s instructions, with the addition of a 0.45 µm syringe filter step (76479-008, VWR International, PA, USA) to remove cells from samples being measured. A Spectramax i3x plate reader (Molecular Devices, CA, USA) was used to quantify the luminescence signal in opaque white 96 well plates (CA25382-208, Corning, NY, USA). Collected samples were quantified for eATP immediately post-collection (**Figure 1B**) or combined with the ATP stabilization buffer and archived at −80°C for up to 6 weeks before quantification (**Figure 1C**).

**Figure 1.**
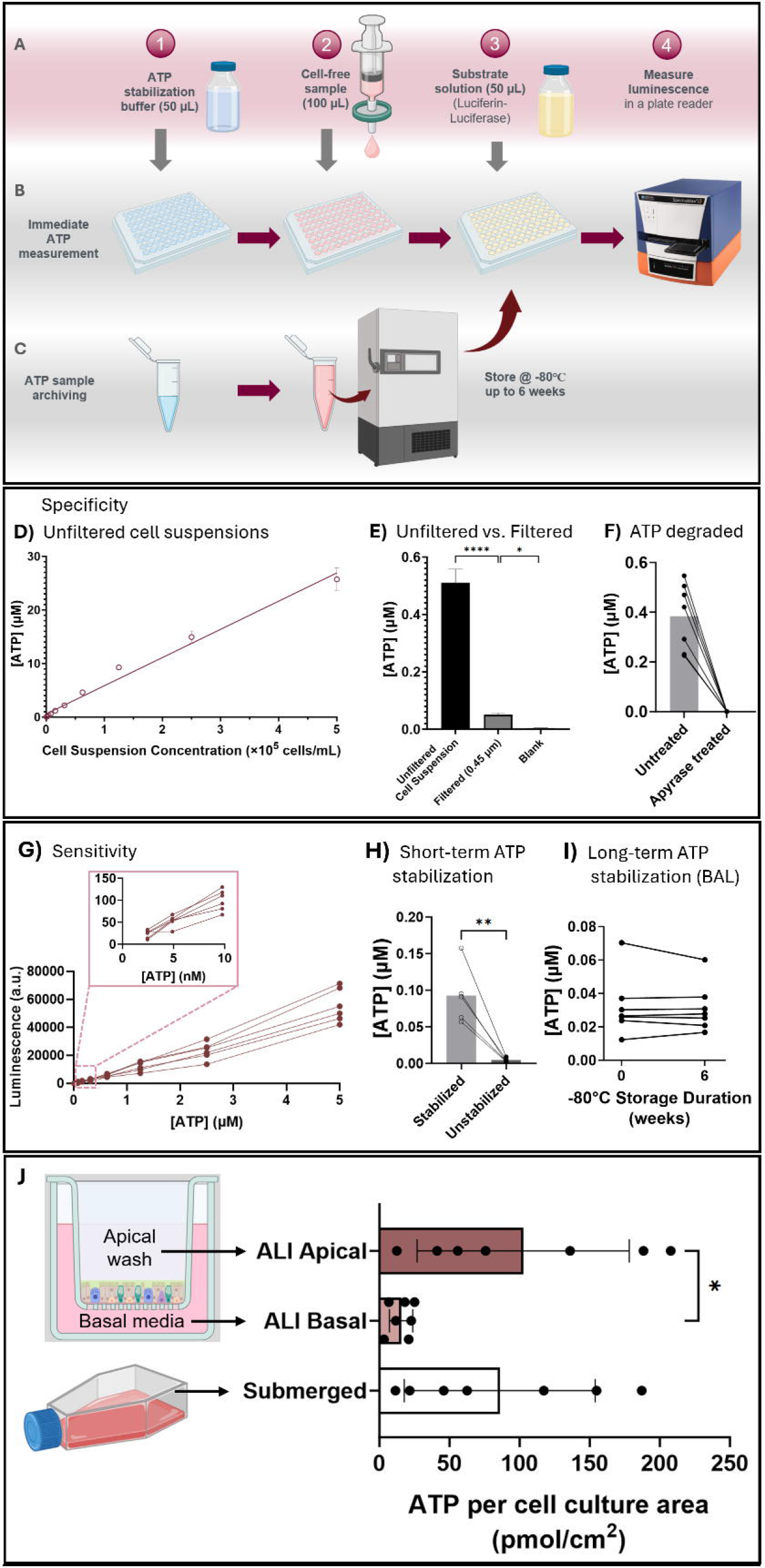
ATPlite™ assay procedure for measuring extracellular ATP (eATP). **A)** Schematic of the general assay procedure for 96 well plates. Cells must be removed from samples using syringe filters to isolate the eATP signal. ATP can be measured from samples **B)** immediately or **C)** archived at −80°C for up to 6 weeks. **D)** ATP concentrations of Calu-3 cell suspensions lysed by the ATPlite™ kit. **E)** ATP concentrations of unfiltered and syringe filtered Calu-3 cell suspensions and blank cell culture media as a negative control. **F)** ATP concentration before and after treatment with apyrase to degrade ATP. **G)** Luminescence intensities of 6 independent ATP standard curves from 0.05 nM to 5 µM. **H)** ATP concentrations of cell culture media samples after 4 h of incubation with ATP stabilization buffer or PBS control. **I)** ATP concentration of bronchoalveolar lavage (BAL) samples before and after 6 weeks of storage at −80°C with ATP stabilization buffer. **J)** Baseline eATP levels from normal human bronchial epithelial cells under air-liquid interface (ALI) and submerged culture conditions. Each plotted point represents individual NHBE donors (N=7). * P < 0.05.

The relationship between the number of cells in a sample and the measured ATP signal was demonstrated by measuring ATP in a series of suspensions of Calu-3 cells, a human lung adenocarcinoma cell line derived from a 25-year-old male (**Figure 1D**). To investigate the ability of 0.45 µm syringe filters to remove cells and a source of measurement noise, ATP was measured from Calu-3 cell suspensions (8000 cells/mL) before and after syringe filtering (**Figure 1E**). The specificity of the assay for ATP was investigated by quantifying ATP in syringe-filtered Calu-3 cell culture media samples before and after treatment with 5 units/mL apyrase, which hydrolyzes ATP into ADP and AMP. (**Figure 1F**). The ATP detection sensitivity was investigated in serial dilutions of ATP standard (Revvity, MA, USA) from 0.05 nM to 5 µM (**Figure 1G**). To evaluate the stability of ATP in cell culture media samples with and without addition of the ATPlite™ ATP stabilization buffer, ATP was measured in syringe-filtered Calu-3 cell culture media samples before and after 4 h of incubation at 4°C with ATP stabilization buffer or PBS (unstabilized) as vehicle control (**Figure 1H**). To evaluate the stability of ATP in human BAL fluid samples after long-term storage, ATP was measured immediately upon sample receipt and after 6 weeks of storage at −80°C with ATP stabilization buffer (**Figure 1I**).

### Quantifying eATP from primary human bronchial epithelial cell cultures

Primary normal human bronchial epithelial (NHBE) cells (Lonza Bioscience) were cultured under submerged conditions in T25 flasks (Eppendorf Canada Ltd., ON, Canada) or ALI conditions in Transwell™ inserts (3470, Corning, NY, USA). In submerged cultures of 80% confluence, eATP was measured from 0.45 µm syringe filtered cell culture media sampled after 2 days of culture (**Figure 1J**). In 3-week differentiated air-liquid-interface (ALI) cell cultures, eATP was measured from basal media samples after 2 days of culture or syringe filtered apical washes with 200 µL of Hank’s Balanced Salt Solution (HBSS; 311-512 CL, Wisent Inc., QC, Canada).Measured ATP concentrations were normalized to the cell culture surface area and media volume for direct comparison between different culture vessels.

## Results

The ATP concentration was directly proportional to the number of Calu-3 cells in suspension, confirming that the ATPlite™ kit releases intracellular ATP, and highlighting the need to remove cells from samples in which measurement of eATP is desired (**Figure 1D**). Using 0.45 µm syringe filters to remove Calu-3 cells from suspension significantly reduced measured ATP levels compared to unfiltered suspensions (**Figure 1E**). This resulting eATP concentration from filtered suspensions was significantly higher than blank cell culture media vehicle controls, demonstrating that filtering is able to isolate eATP signal. Addition of apyrase to Calu-3 cell culture media samples abolished all luminescence (**Figure 1F**), showing that the ATPlite™ signal is ATP-specific and is not responsive to its metabolites ADP or AMP. The luminescence signal provided by the ATPlite™ assay was linear from 5nM to 5µM ATP, below which the measurements were not different from blank controls (**Figure 1G**). Syringe-filtered cell culture media samples treated with ATP stabilization buffer maintained their ATP concentrations after 4 h of incubation at 4°C, whereas those treated with only vehicle had diminished ATP concentrations (**Figure 1H**). BAL samples treated with ATP stabilization buffer maintained ATP concentrations following 6 weeks of storage at −80°C relative to paired samples measured on the day of collection (no freezing), supporting the ability for bulk measurement of multiple samples collected over an extended study duration (**Figure 1I**).

ATP was detectable in cell culture media samples of NHBE submerged monolayer cell cultures (**Figure 1J**). In ALI cultures, ATP was predominantly released on the apical side compared to the basal compartment. Submerged monolayer cultures had eATP levels comparable to that of the apical surface of ALI cultures.

## Conclusion

Our methods and data describe measurement of ATP with filtration using commercially available products, for isolation of eATP signal and removal of intracellular ATP. The approach results in higher ATP signal stability than typical transient luminescence microwell plate-based assays. The measured eATP concentrations from primary human bronchial epithelial cells were consistent with those observed in studies using real-time luminescence measurement.[12, 13] Our method revealed that ATP is preferentially secreted on the apical lumen of human airway epithelial cells at rest. This apical preference is logical given that basal ATP release in the absence of stimulation is mainly driven by pannexin channels which are expressed apically in ALI-differentiated ciliated bronchial epithelial cells.[14, 15]. Lastly, we demonstrate the stability of ATP in frozen BAL samples for at least 6 weeks, allowing samples to be stored and measured in larger batches to reduce batch-batch variability between experiments. Our results and approach are directly relevant to the pre-clinical and clinical research exploring P2X3 receptor antagonists and experimental interventions for cough and other airway diseases.

## Notes

**Conflict of interest**: RS, EK, QC, JPN, WH, and SB, have no conflicts of interest to disclose relevant to this manuscript. J.A.H. reports grants from Merck (for this study) and consulting fees from GSK outside the submitted work. **J.A.H.** is founder, equity holder, and CSO of Tessella Biosciences which operates outside of the submitted work. **I.S.** is supported by the Canada Research Chair program, reports grants from Merck (for this study), MITACS, GSK, Bellus, Trevi Therapeutics, Bayer, Genentech; personal speaker fees from Merck, GSK, AstraZeneca, Sanofi-Regeneron, Nocion; consulting fees from Merck, GSK, Bellus, Sanofi-Regeneron, Methapharm outside the submitted work.

**Funding**: Supported in part by a research grant from Investigator-Initiated Studies Program of Merck Canada Inc. The opinions expressed in this paper are those of the authors and do not necessarily represent those of Merck Canada Inc.

### Competing Interest Statement

J.A.H. reports grants from Merck (for this study) and consulting fees from GSK outside the submitted work. J.A.H. is founder, equity holder, and CSO of Tessella Biosciences which operates outside of the submitted work. I.S. is supported by the Canada Research Chair program, reports grants from Merck (for this study), MITACS, GSK, Bellus, Trevi Therapeutics, Bayer, Genentech; personal speaker fees from Merck, GSK, AstraZeneca, Sanofi-Regeneron, Nocion; consulting fees from Merck, GSK, Bellus, Sanofi-Regeneron, Methapharm outside the submitted work.

